# Unraveling the substrate preference of an uncharacterized phylogenetic subgroup in Formate/Nitrite Transporter (FNT) family: Computational studies of anion transport in *Escherichia coli* FNT homolog

**DOI:** 10.1101/637116

**Authors:** Mishtu Mukherjee, Ramasubbu Sankararamakrishnan

**Affiliations:** Department of Biological Sciences and Bioengineering, Indian Institute of Technology Kanpur, Kanpur, India

## Abstract

Formate/Nitrite Transporters (FNTs) selectively transport monovalent anions and are found in prokaryotes and lowers eukaryotes. They play significant role in bacterial growth and act against the defense mechanism of infected host. Since FNTs don’t occur in higher animals, they are attractive drug targets for many bacterial diseases. Phylogenetic analysis revealed that they can be classified into eight subgroups and two of which belong to the uncharacterized YfdC-α and YfdC-β groups. Experimentally determined structures of FNTs belonging to different phylogenetic groups adopt the unique aquaporin-like hourglass helical fold. We considered formate channel from *Vibrio Cholerae* (VcFocA), hydrosulphide channel from *Clostridium difficile* (CdHSC) and the uncharacterized channel from *Escherchia coli* (EcYfdC) to investigate the mechanism of transport and selectivity. Using equilibrium molecular dynamics (MD) and umbrella sampling studies, we determined temporal channel radius profiles, permeation events and potential of mean force (PMF) profiles of different substrates with the conserved central histidine residue in protonated or neutral form. Unlike the VcFocA and CdHSC, MD studies showed that the formate substrate was unable to enter the vestibule region of EcYfdC. Absence of a conserved basic residue and presence of acidic residues in the vestibule regions, conserved only in YfdC-α, were found to be responsible for high energy barriers for the anions to enter EcYfdC. PMF profiles generated for ammonia and ammonium ion revealed that EcYfdC can transport neutral solutes and could possibly be involved in the transport of cations analogous to the mechanism proposed for ammonium transporters. Although YfdC members belong to the FNT family, our studies convincingly reveal that EcYfdC is not an anion channel. Absence/presence of specific charged residues at particular positions makes EcYfdC selective for neutral or possibly cationic substrates. This adds to the repertoire of membrane proteins that use the same fold but transport substrates with different chemical nature.

**Author Summary:** Channels and transporters are membrane proteins involved in the transport of solutes selectively across the cell membranes. Drugs for many diseases have been developed to inhibit ion channels. Formate/Nitrite Transporters (FNTs) are ion channels selective for monovalent anions and are present in bacteria and lower eukaryotes. Absence of FNTs in humans makes them as attractive drug targets against many pathogenic bacteria. To develop inhibitors for a protein, it is important to understand the mechanism of its function. Selectivity and transport mechanism of FNTs have been investigated for some members. One of the subgroups of FNTs, YfdC-α, is uncharacterized. In this study we used computer simulation approach to investigate the molecular mechanism of selectivity and transport of three FNTs including one from YfdC-α group from *Escherichia coli*. Our studies show that *E. coli* YfdC is not an anion channel although it belongs to FNT family. We hypothesize that the YfdC-α members could be involved in the transport of neutral or possibly cationic substrates. This is further supported by the conservation of specific acidic residues found only in YfdC-α in the vestibule regions. This finding has major implications in developing blockers for FNT members belonging to YfdC-α group.

## Introduction

The family of Formate/Nitrite Transporters (FNTs) is selective for monovalent anions and members of this family are involved in the transport of anions such as formate, nitrite, hydrogen sulphide and lactate [1–8]. They are present in bacteria, archaea and lower eukaryotes and are not identified in mammals including humans [9]. The formate channel FocA, the nitrite channel NirC and the hydrosulphide channel HSC are the prototype members of the ancient FNT family [6,7,10]. FNT members facilitate bacterial growth under anaerobic conditions, enable the bacteria to survive against the defense mechanism of the infected host and serves as an essential factor of the energetic flux in malarial parasites. Three-dimensional structures of few FNT family members representing different subgroups have been determined. This includes formate-specific FocA [11,12], nitrite-transporting NirC [4] and hydrogen sulphide-selective HSC channels [7]. All FNT members from different sources with different substrate selectivity adopt the same helical fold with six transmembrane helices (TM1 to TM6). TM2 and TM5 are interrupted in the middle of the transmembrane region giving rise to two fragments. Ω-loop connects TM2a and TM2b while S-loop links TM5a and TM5b. The fold of FNT members is strikingly similar to the hourglass helical fold formed by Major Intrinsic Protein (MIP) channels of which aquaporins and aquaglyceroporins are the two prominent members [13]. However, unlike MIP channels which form tetramers, FNT members form pentamers under physiological conditions. The selectivity and transport mechanism of FNT members have been investigated by both experimental and computational techniques.

Hunger et al. [14] developed a reporter assay to monitor the changes in the formate level in cytoplasm of *Escherichia coli*. They studied variants of the formate channel, FocA. Residues at the central constriction site, cytoplasmic slit, Ω-loop and the N-terminal helix were substituted at 16 different positions. Their studies revealed the role of the conserved residues in regulating the direction of formate transport and possible role in gating of FocA channel. In another study Doreen et al. [15] identified a three-residue C-terminal motif in FocA that is important for formate passage. Molecular dynamics (MD) simulations of FNT family members have been carried out to characterize the protein’s function and to resolve the question whether this protein functions as a channel or transporter. Feng et al. [16] performed MD simulations on the crystal structures of two FocA structures determined from *Salmonella typhimurium* (StFocA) [17] and *E. coli* (EcFocA) [11]. Both neutral and protonated states were considered for the highly conserved His 209 (residue numbering corresponds to PDB ID: 3KCU [11]) to mimic the high and low pH conditions. In this study, the role of N-terminal helix and the interactions of neutral/protonated His-209 have been highlighted in the opening and closing of the channels. However, the simulation time of 20 ns may not be sufficient and later MD simulation studies showed that N-terminal helix of each protomer does not occlude the channel and the substrate can permeate crossing the N-terminal helix [18]. Simulation studies on *Vibrio cholerae* FocA (VcFocA) [12] also offered explanation for the diminished electrical current observed in electrophysiological experiments in low pH. Lv et al [18] hypothesized that the substrate transported at low pH could be formic acid and the protonation state of His 209 plays a vital role in the transport of formate/formic acid substrates. Atkovska and Hub [19] employed a variety of computational techniques to study the transport mechanism of FNT members. The computed potential of mean force profiles demonstrated that anions cannot permeate the NirC, HSC and FocA proteins irrespective of whether the central His residue is protonated or neutral. Their subsequent studies on NirC pore concluded that anion protonation is required during the permeation which could happen by proton transfer between the central His and the anion substrate. The authors describe FNT channels as “the protein family existing on the border between channels and transporters”. Recently, Wiechert and Beitz [20] developed direct transport assays using radiolabeled substrates and reported the role of a conserved lysine residue in the periplasmic vestibule in the transport of anions. By combining mutagenesis and computational studies, they demonstrated increased transport of neutralized substrate in acidic pH and proposed that this substrate neutralization happens in the bulk water region close to the vestibule, thus leading to dielectric shift of substrate acidity. They drew parallel to the transport mechanism of ammonium transporters [21,22] and suggested that the models explaining the transport of weak acid and base neutralizing transporters are complementary to each other.

In addition to published reports related to the structure and transport mechanism of FNTs, it is important to understand the extent of diversity within FNT family and such studies will provide knowledge regarding the evolution of FNTs and the diverse solutes that are likely to be transported selectively by FNT members. We have recently carried out extensive bioinformatics analysis of more than 2200 FNT members from bacteria, archaea and eukaryotes [9]. Structure-based sequence alignment revealed high conservation of nearly one third of all residues within the transmembrane region. Eight different subfamilies have been identified using phylogenetic analysis. Apart from the known subfamilies that include formate, nitrite and hydrosulphide transporters, two subfamilies, YfdC-α and YfdC-β, with unassigned function were also recognized. Both taxonomic distribution and sequence conservation at specific positions exhibit subfamily-specific features. In this study, we have performed equilibrium molecular dynamics simulations on representative FNT members (FocA, HSC and YfdC) and showed that the formate anion substrate molecules cannot even enter the periplasmic and cytoplasmic vestibule regions of the *E. coli* YfdC channel. Free energy profiles of the substrates derived using umbrella sampling studies indicate that the *E. coli* YfdC channel is not an anion transporter and is likely to transport neutral solutes or even cations. Our simulation studies are further supported by the subfamily-specific conservation of residues found at the vestibule regions of YfdC subfamily.

## Results and Discussion

We first performed equilibrium molecular dynamics simulations on three representative FNT members, namely VcFocA, CdHSC (FNT homolog from *Clostridium difficile* selective for hydrosulphide) and EcYfdC. Each FNT member was simulated in the presence of the substrates formate or formic acid and we have considered the highly invariant central His residue (His 209 in PDB ID: 3KCU numbering) in either protonated or neutral state (Table 1). Thus a total of nine simulations were performed each for a period of 500 ns as described in the Methods section. We first analyzed the overall channel behavior of all the three channels by plotting the temporal channel radius profiles (Fig 1). In all the systems, the cytoplasmic slit and the central constriction regions were narrow and the channel radius less than 1.4 Ǻ was maintained throughout the simulations in these constriction regions. Among the three channels, VcFocA showed the most relaxed pore (Fig 1A, Fig 1D and Fig 1G) and this can be understood from the fact that the Ω-loop and TM2b segment are flexible regions. The EcYfdC channel exhibited the narrowest pore compared to the other two FNT subtypes (Fig 1C, Fig 1F and Fig 1I). The behavior of HSC channel was in between that of VcFocA and EcYfdC (Fig 1B, Fig 1E, Fig 1H). When the central His was neutral and the substrate was formic acid, all three systems remained tight in both constriction regions (Fig 1G, Fig 1H, Fig 1I) and this was especially true for EcYfdC in which the pore was closed from the cytoplasmic to extracellular side (Fig 1I). In general, relaxation of the channel was observed when the central His was protonated and the permeating substrate was either formic acid or formate ion. It must be noted that none of the nine systems exhibited fully open states due to the fact that both constriction regions always remained closed in all the conditions we tested. Previous studies have mentioned that FNTs are slow channels or substrate passage is either gradient-dependent or regulated [4,7,18,23].

**Table 1:** Summary of all the equilibrium MD simulations performed on the FNT systems and the number of permeation events for each system

| FNT system<sup>a</sup> | Protonation state of the Central His<sup>b</sup> | Solute (120 mM)<sup>c</sup> | Number of permeation events<sup>d</sup> |  |
| --- | --- | --- | --- | --- |
|  |  |  | Formic acid | Water |
| VcFocA | Protonated | Formate | N/A | 109 |
|  | Protonated | Formic acid | 10 | 110 |
|  | Neutral | Formic acid | 4 | 0 |
| CdHSC | Protonated | Formate | N/A | 37 |
|  | Protonated | Formic acid | 2 | 52 |
|  | Neutral | Formic acid | 1 | 0 |
| EcYfdC | Protonated | Formate | N/A | 16 |
|  | Protonated | Formic acid | 1 | 23 |
|  | Neutral | Formic acid | 0 | 0 |

<sup>a</sup>For the formate channel VcFocA from *V. cholera* and the hydrosulphide channel CdHSC from *C. difficile*, the starting structures correspond to the PDB Ids 3KCU and 3TE0 respectively. For the EcYfdC channel from *E. coli*, we generated homology models as described in the main text. All simulations were performed for a period of 500 ns production run.
<sup>b</sup>Central His (His 209 in 3KCU numbering) is the histidine residue that is invariant across all FNT members and forms part of the central constriction. This His in neutral form was considered only when formic acid was considered as substrate. For formate ion, central His in neutral state would be energetically unfavorable.
<sup>c</sup>Saturating concentrations of formic acid/formate were used as reported in [ref]. Simulations under equilibrium conditions were performed in the absence of any solute gradient.
<sup>d</sup>A permeation event is said to occur if the substrate completely traverses through a cylinder of 0.5 nm passing through each monomer along the channel axis. N/A: Not Applicable

**Fig 1:**
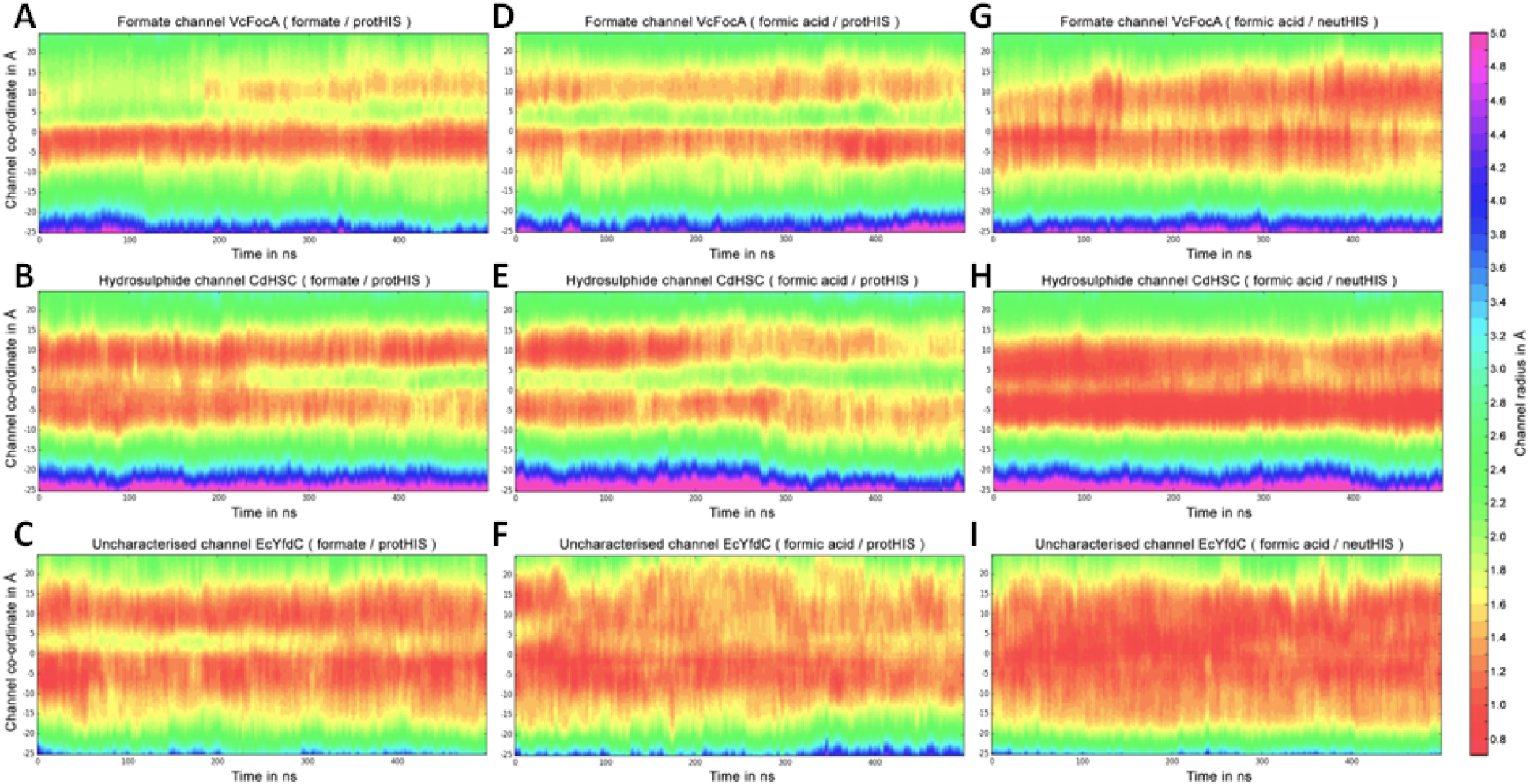
Temporal channel radius profiles of all the nine simulated systems. The temporal channel radius profiles are plotted for 500 ns simulation time for VcFocA, CdHSC and EcYfdC channels. (A-C) Central His is protonated and the substrate is formate ion; (D-F) Central His is protonated and the substrate is formic acid; (G-I) Central His is neutral and the substrate is formic acid. Position ‘0’ in the Y-axis corresponds to the central His residue. The positive and negative Y-axis values correspond to periplasmic and cytoplasmic sides respectively. HOLE [24] profile data from all five monomers in each time frame have been combined to create the time evolution of channel radius profiles. See Methods section for further details.

### Conformation of central His residue in FNT channels: protonated vs neutral

Earlier simulation studies of VcFocA channel indicated that central His residue can adopt two conformational states namely, downward and upward states. We analyzed the His-209 conformations in all the nine simulations. The distribution of side-chain dihedral angles of central His residue is shown in Fig 2. We observed that whenever the central His was protonated, it mostly preferred to assume upward state and this corresponds to its side-chain dihedral angle χ_1_ adopting ± 45°. This was in contrast to what we saw in simulations in which the central His was considered as neutral. The χ_1_ of central His preferred ±180° in the neutral form which corresponds to the downward state. In this state, the hydrogen bond between the central His and the Thr residue (Thr 91 in 3KCU numbering) remained stable throughout the simulations (Data not shown). The preference for His-209 to adopt upward state when it was protonated seemed to be responsible for increased solvation of the channel and increased fluctuation of another central constriction site residue, Phe- 202 from TM2b. Both factors contributed to the relatively wider pore compared to the simulations in which His-209 was considered neutral.

**Fig 2:**
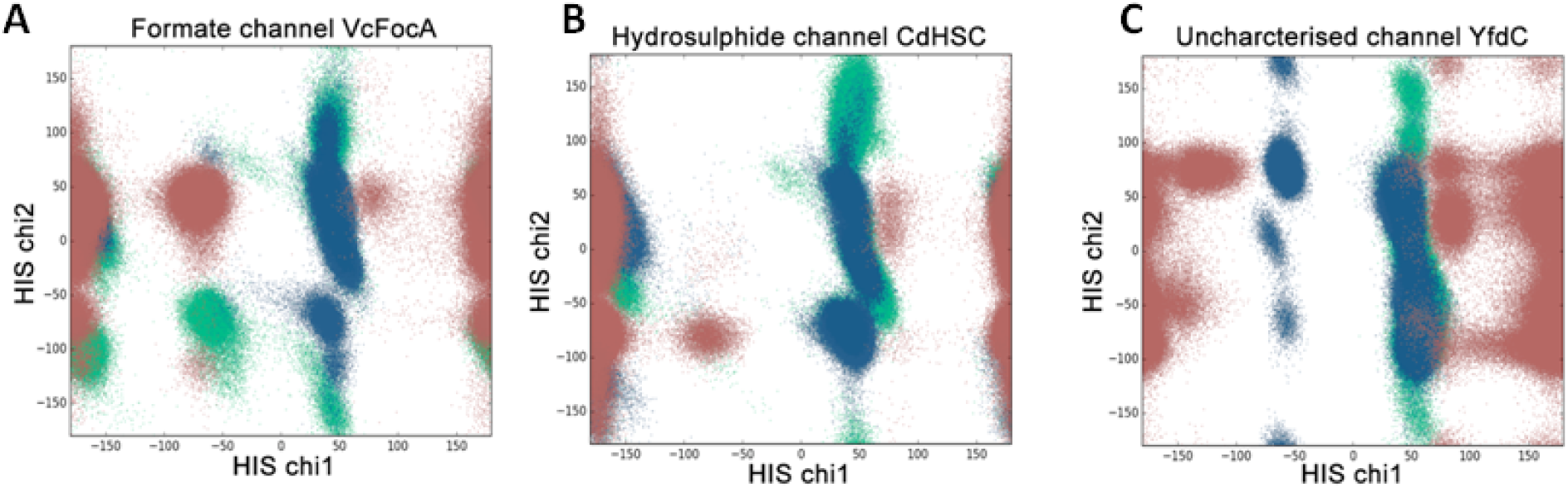
Distribution of χ_1_ vs χ_2_ side-chain dihedral angles of the central His residue. The χ_1_-χ_2_ angles for the central His residue are plotted for (A) VcFocA, (B) CdHSC and (C) EcYfdC systems. Depending upon the substrate and the protonation state of the central His, the data points are shown in different colors. Green: substrate-formate and central His-protonated; Blue: substrate-formic acid and central His- protonated; Pink: substrate-formic acid and central His-neutral.

## Permeability of solutes in the simulated FNT systems

We examined the number of permeation events in all three FNT systems for both formate ion and formic acid. A permeation event was defined if the substrate completely traversed the transmembrane region from one side of the membrane to the other side. For each system, we plotted the Z-coordinate of the center of mass of formate ion or formic acid as a function of time along the channel coordinate. MD trajectories of Z-coordinates of center of mass of individual substrates along the channel coordinates are plotted in Fig 3.

**Fig 3:**
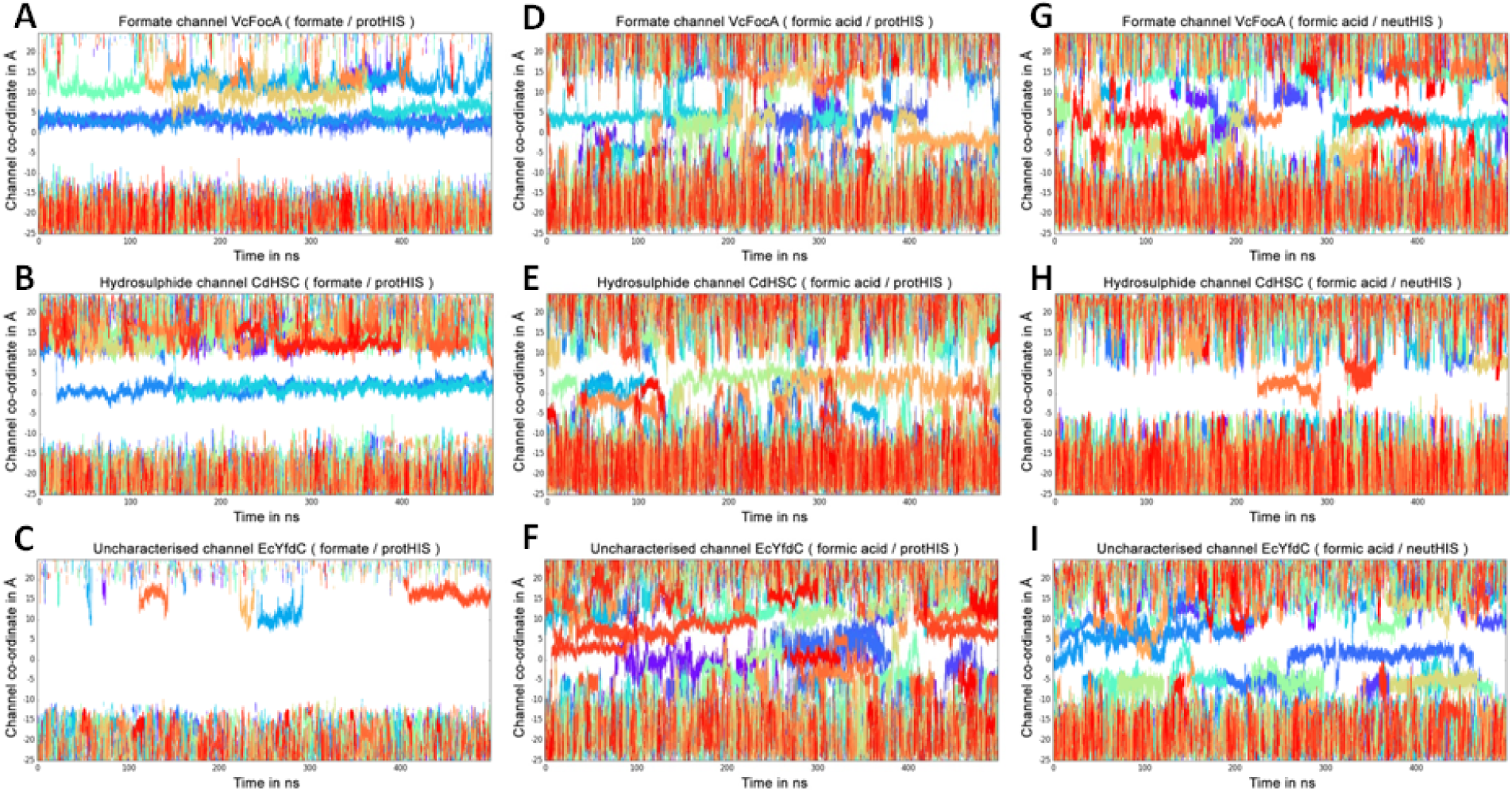
MD trajectories of the position of the substrate (formate ion or formic acid) along the channel coordinate. Evolution of Z-coordinates of the center of mass of individual substrate molecules as a function of time are plotted for the entire 500 ns production run for all nine systems. Each color represents an individual substrate molecule. Analysis is shown for VcFocA, CdHSC and EcYfdC systems. (A-C) substrate-formate ion and central His- protonated; (D-F) substrate-formic acid and central His-protonated; (G-I) substrate-formic acid and central His-neutral. Position ‘0’ in the Y-axis corresponds to the central His residue. The positive and negative Y-axis values correspond to periplasmic and cytoplasmic sides respectively.

First we looked at the behavior of formate ions in all the FNT systems studied when the central His was protonated. Several formate ions entered VcFocA and CdHSC channels. However when the central His residue remained in the protonated state, they were not able to go beyond this residue (Fig 3A and Fig 3B). Earlier simulation studies also observed a similar phenomenon in the case of formate and nitrite channels [18,19]. The calculated PMF profiles showed an energy minimum for the anionic species when His was protonated in these channels. It was suggested that presence of two formate ions in the channel could help in reducing the energy barrier. Alternatively, it was hypothesized that a proton transfer from the central His could enable the completion of permeation process [18,19]. In our analysis, we found that presence of another nearby formate ion did not produce any significant change and the substrate continued to reside close to the central His. When all three FNT members are compared, we see a distinct difference in EcYfdC channel. Only a couple of formate ions entered the vestibule regions of EcYfdC channels (Fig 3C). While entry of formate ions up to the protonated central His was frequent in the case of VcFocA and CdHSC channels, there was not a single instance of formate ion entering the vestibular regions and approaching the central His residue in the case of EcYfdC channel. This is one of the most notable features that distinguish EcYfdC from the other two FNT members.

When the substrate was formic acid, the permeation behavior was different in all the three FNT systems. Being neutral, formic acid entered both cytoplasmic and extracellular vestibules in all the FNT subtypes and reached the central His irrespective of its protonation state (Fig 3D to Fig 3I). We also observed occasional diffusion through the constriction regions resulting in a complete permeation. VcFocA with the protonated central His and formic acid as the substrate exhibited a maximum of 10 permeation events during the 500 ns simulation (Table 1). This is followed by VcFocA with neutral His showing 4 full permeation events. Both CdHSC and EdYfdC showed permeation events rarely, only one or two during the entire course of simulation. This could be explained by the fact that the cytoplasmic slit region was tightly closed during most of the simulations in these two channels (Fig 1). Permeation events reported in this study mostly agree with the earlier non-equilibrium MD simulations on FocA channels [18,19].

## Water permeation in FNT systems

In the next step, we have examined the permeation events of water molecules across the FNT channels in all the nine simulations (Table 1). When central His was protonated, we observed that on an average VcFocA permeated more than 100 water molecules through its monomers when the simulation was performed with formate or formic acid. Although CdHSC exhibited water permeation events, its monomers transported relatively less number of water molecules compared to VcFocA. The average number of water molecules translocated through CdHSC system is between 38 to 50 while this number is even less for EcYfdC channel. As few as 16 to 23 water molecules were transported through EcYfdC monomers indicating that the constriction regions are the narrowest in this system. Interestingly, when central His was neutral, not a single permeation event occurred in all the three simulated systems irrespective of the substrates considered. The average number of water molecules even for VcFocA was still lower than those conducted by a typical water-transporting aquaporins [25]. Hence, one could consider the water permeation in FNT systems as “leakage” instead of true permeation events. This observation is consistent with the stop-flow experiments which showed minimal water transport in FocA channels [11].

Presence of a water file inside FNT channels will give rise to the possibility of proton transport by Grotthuss mechanism whereby the protons utilize the continuous hydrogen bonded chain in the water file to traverse the transmembrane region. To establish or to rule out the possibility of a proton transport, we calculated the average dipole orientation (<cosθ>) of all water molecules along the channel axis at different z-coordinates. Here θ is the angle between the water dipole vector and the membrane normal. We performed this analysis for both VcFocA and CdHSC systems and plotted the water dipole orientation profiles (Fig 4). We observed an abrupt switch in the dipole orientation in the middle of the channel near the central His residue indicating that this His is likely to play a role similar to that found in the Asn of NPA motif in aquaporin channels to prevent proton leakage [26–28].

**Fig 4:**
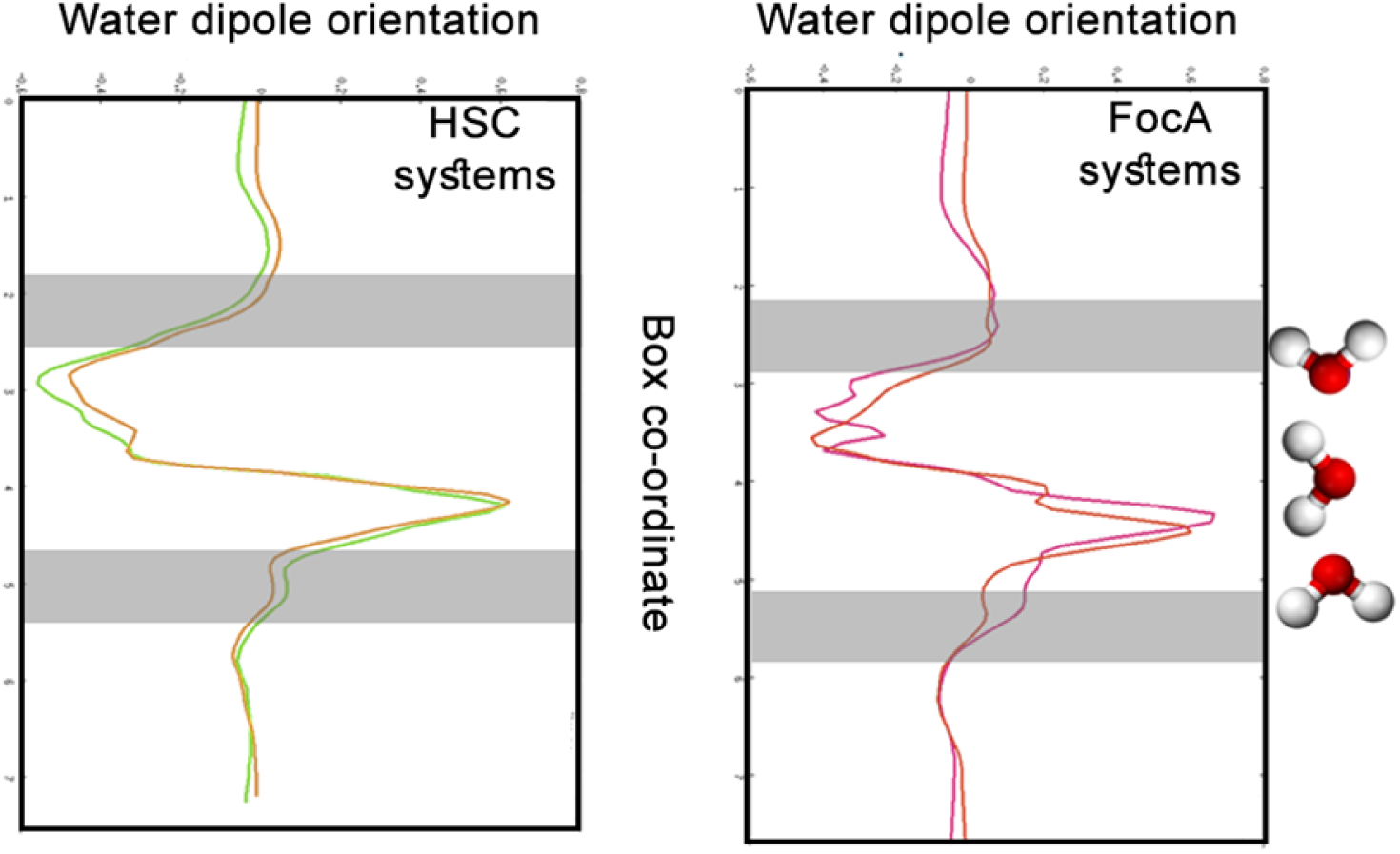
Dipole orientation of water molecules inside the VcFocA and CdHSC systems. Dipole orientation of water molecules inside the channel is shown for CdHSC and VcFocA systems. The analysis is shown for the systems in which the central His is protonated. Dipole flip near the center of the channel is evident. The gray bands mark the lipid head-group region. Water molecules are plotted to show the change in overall dipole orientation.

The above analyses show that the behavior of EcYfdC with the unassigned function is distinct in terms of water conduction and channel characteristics. YfdC members constitute nearly one fourth of all FNT family members [9]. Our previous bioinformatics analysis showed that some key positions in YfdC members are substituted although they retain overall characteristics of FNT family [9]. MD simulation studies clearly demonstrated that EcYfdC channel is not efficient in attracting anionic formate ion in its vestibular regions. This is in stark contrast to the VcFocA and CdHSC channels where formate ions went beyond the vestibular regions and reached the middle of the channels. However although no full permeation was observed, the extent of permeation of neutral substrate, formic acid, was comparable in all the three channels simulated. Hence, we suspected that YfdC channels may not be amenable for anion transport. To test this hypothesis, we designed the following computational experiment on EcYfdC channel and the results of the computational studies are presented in the subsequent sections.

## Role of periplasmic helix in substrate permeation

In all the three FNT channel simulations, one of the major differences is the behavior of formate ions at the vestibule regions. While formate ions entered both the vestibules in VcFocA and CdHSC systems, this substrate could not enter the periplasmic vestibule of EcYfdC (Fig 3). However, this was not the case with the formic acid and this neutral substrate entered both the vestibule regions and the behavior of the substrate at the channel entrance and in the interior was very similar in all three FNT members. Hence, we examined the possible factors that distinguish YfdC family from other FNT family members to find why anionic substrates could not enter the vestibule regions. We found a highly conserved Lys residue (Lys-156 in PDB ID: 3KCU numbering) in the so called periplasmic helix (P-helix) at the periplasmic side. This helix connects the two halves of the channel linking TM3 and TM4 helices and Lys-156 is located near the C-terminal end of the P-helix with its side-chain pointing towards the interior of the channel (Fig 5A). This residue has been shown to be experimentally important for formate permeation [20] and previous simulation studies suggest that this periplasmic Lys facilitates the entry of the formate substrate [18]. Sequence analysis of residues in the P-helix region exhibits near absolute conservation of Lys-156 in formate channels and hydrosulphide channels (Fig 5C). The equivalent position in YfdC-α subgroup lacks a Lys residue. However, we noticed that there is a Lys/His residue in the preceding position in EcYfdC and some other YfdC-α members. When we modeled EcYfdC, we manually adjusted target-template sequence alignment produced by the Salign tool of MODELLER so that this Lys in the model occupied the same position as that of Lys-156 in other FNT members. This model was used in our simulations.

**Fig 5:**
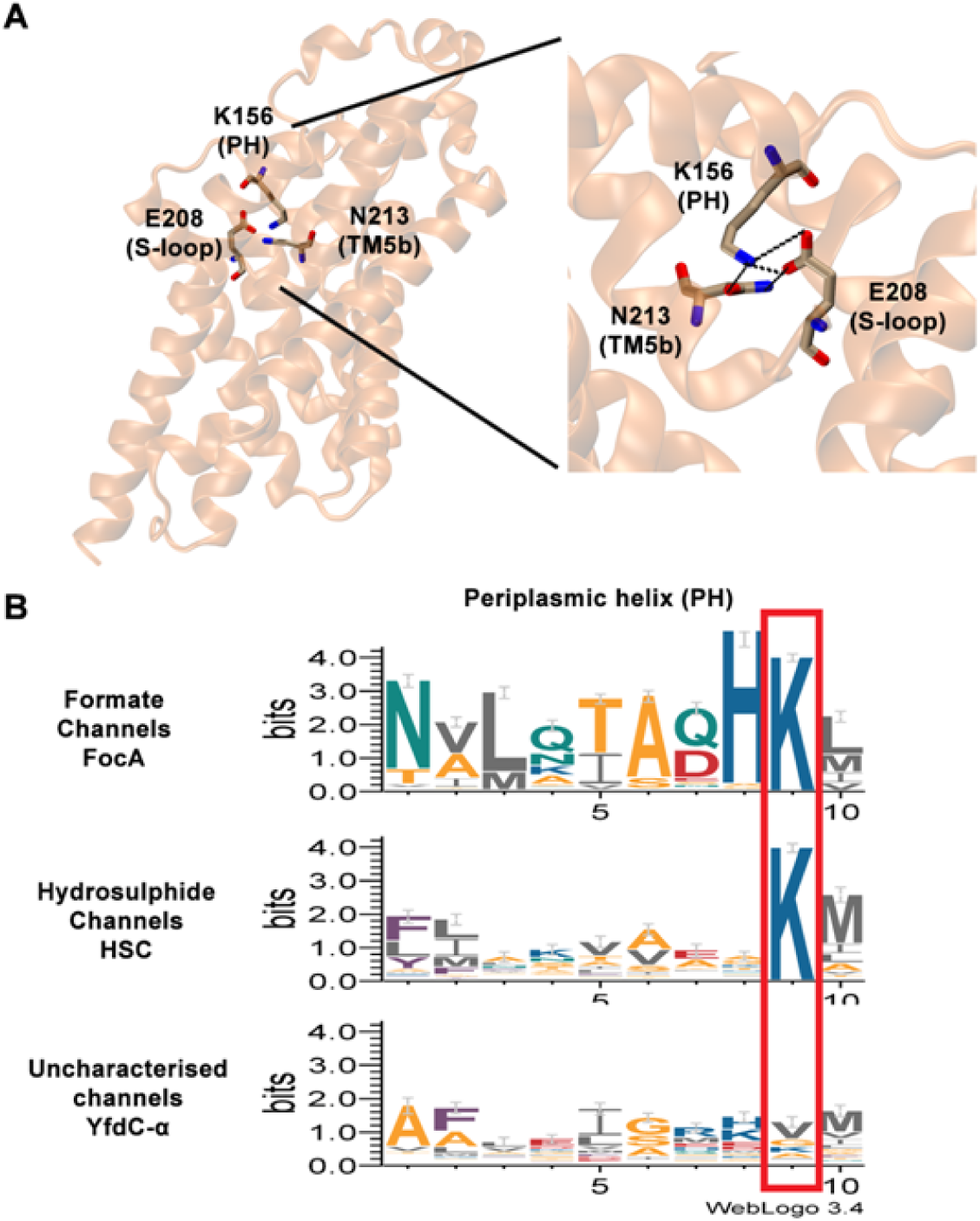
Location and conservation of Lys-156 residue and its interaction with neighboring residues. (A) Formate channel structure (PDB ID: 3KCU) showing the location of Lys-156 residue in the periplasmic helix and its interacting partners Glu-208 from S-loop and Asn-213 from TM5b helix. (B) Salt-bridge and hydrogen bond interactions of Lys-156 with Glu-208 and Asn-213 residues. (C) Sequence Logo of P-helix region shown for FocA, HSC and YfdC-α subfamilies. Position of Lys-156 in FocA and HSC subfamilies and the equivalent position in YfdC-α subfamily are highlighted within a red rectangular box. It is clear that Lys-156 exhibits absolute conservation in FocA and HSC subfamilies while the same position in YfdC-α shows enormous variation.

In both VcFocA and CdHSC simulations, formate ions interacted with Lys-156 for a longer duration while no such interactions were observed in the EcYfdC system. We also examined the stability of periplasmic helix in all nine simulations. The P-helix region lost significant helical character in all three EcYfdC simulations (see below). The lysine residue preceding the 156 position (PDB ID: 3KCU numbering) in EcYfdC moved out of the periplasmic vestibule exposing its side chain to the bulk solvent in all three YfdC systems. This seems to be the major cause for EcYfdC for not able to attract formate ions towards the periplasmic vestibular region.

Since Lys-156 in the periplasmic helix is almost invariant in all the FNT families with the exception of YfdC, we examined its interactions with other residues in FocA and HSC crystal structures. We found out that Lys-156 forms a salt-bridge and hydrogen bond respectively with Glu-208 and Asn-213 (numbering correspond to PDB ID: 3KCU) residues (Fig 5B). Glu-208 occurs in S-loop and Asn-213 is part of the TM5b region. We also found out that both residues are highly conserved in all non-YfdC FNT families (Fig 6). EcYfdc belongs to YfdC-α subgroup and these positions are occupied by small residues in YfdC-α. This explains the reasons for the instability of P-helix in EcYfdC. Even though an adjacent Lys residue was manually adjusted to occupy the same position as that of Lys-156, this Lys lacked interaction partners from S-loop and TM5b as observed in other FNT families. This happened due to substitutions of interacting residues at the equivalent positions.

**Fig 6:**
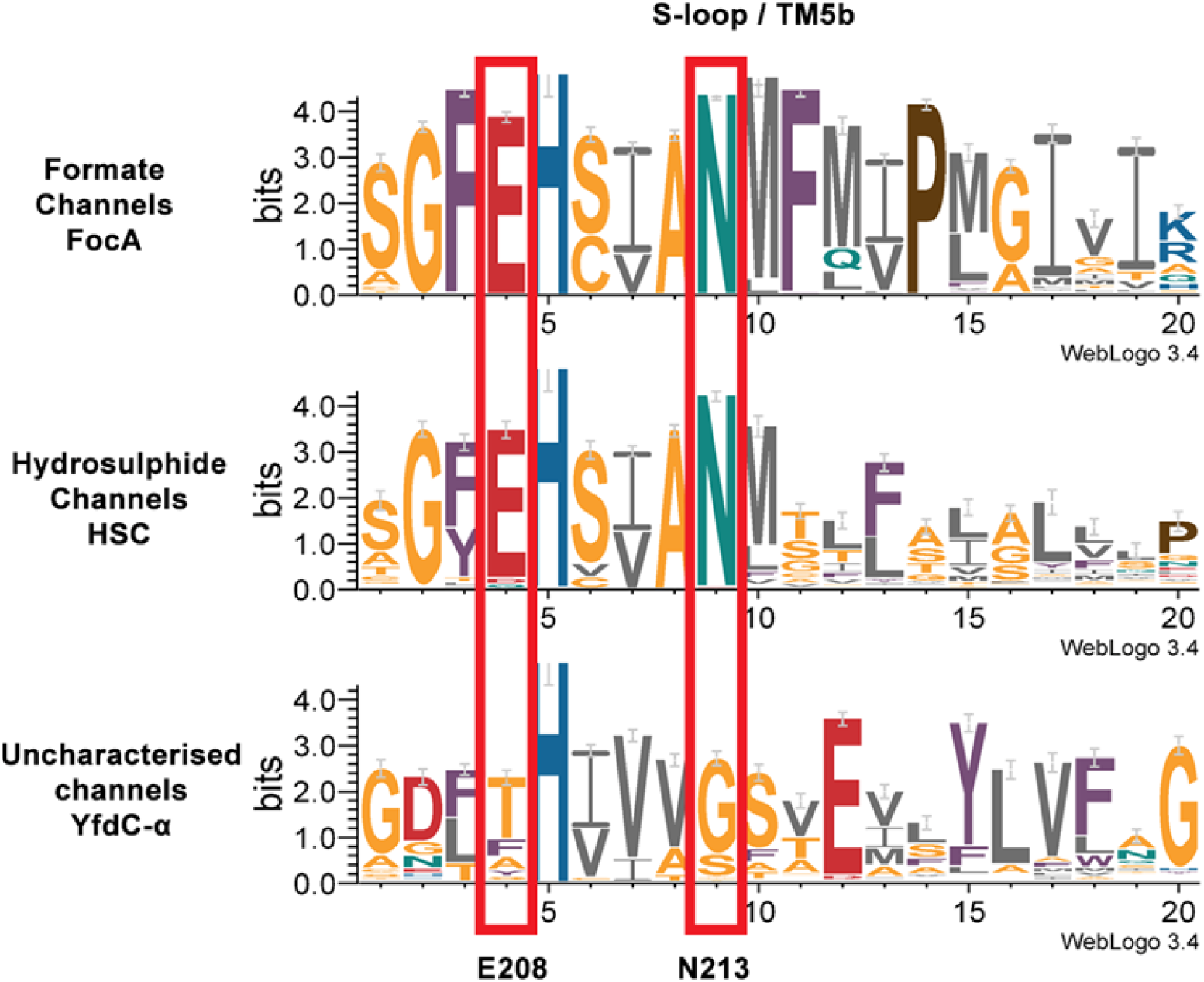
Residue conservation in the S-loop/TM5b region shown for different FNT families. Sequence logo generated for the S-loop/TM5b region for the FocA, HSC and YfdC-α subfamilies. Residues interacting with Lys-156 of P-helix show near absolute conservation in FocA and HSC subgroups and these positions are highlighted within red rectangular boxes.

## MD simulations with revised model of EcYfdC

We reexamined our earlier modeling protocol and decided to use the default alignment produced by the Salign module of MODELLER to generate the homology model of EcYfdC. In the default target-template sequence alignment, a Val residue occupies the position equivalent to Lys-156 in the P-helix region. There was no change in the rest of the model including the transmembrane helical segments and the central His was considered protonated. We performed another 500 ns simulation using the revised model as the starting structure following exactly the same protocol in the presence of 120 mM formate ion. The EcYfdC pentamer at the end of the 500 ns simulation is shown in Fig 7B. It is clear that the helical character of P-helix region remained stable in all five monomers. The Val residue which occupies the equivalent position as that of Lys-156 in other FNT members is facing the vestibular region. The overall behavior of EcYfdC channel in the revised model is similar to that found for the earlier EcYfdC simulation (see above; Fig 7A). This includes features found in the temporal radius profile showing tight constriction regions (S1A Fig). As observed in the previous EcYfdC simulation, entry of formate ions in the perplasmic vestibular regions was not found favorable (S1B Fig) indicating the crucial role played by Lys-156 in facilitating the entry of anionic substrates inside the pore. Hence, we raised the very fundamental question whether YfdC-α members in general and EcYfdC in particular can be considered as anion channels although YfdCs clearly belong to the FNT family [9]. To understand the nature of the substrates permeated by YfdC-α, we calculated the energetics of permeation of various of substrates through EcYfdC channel.

**Fig 7:**
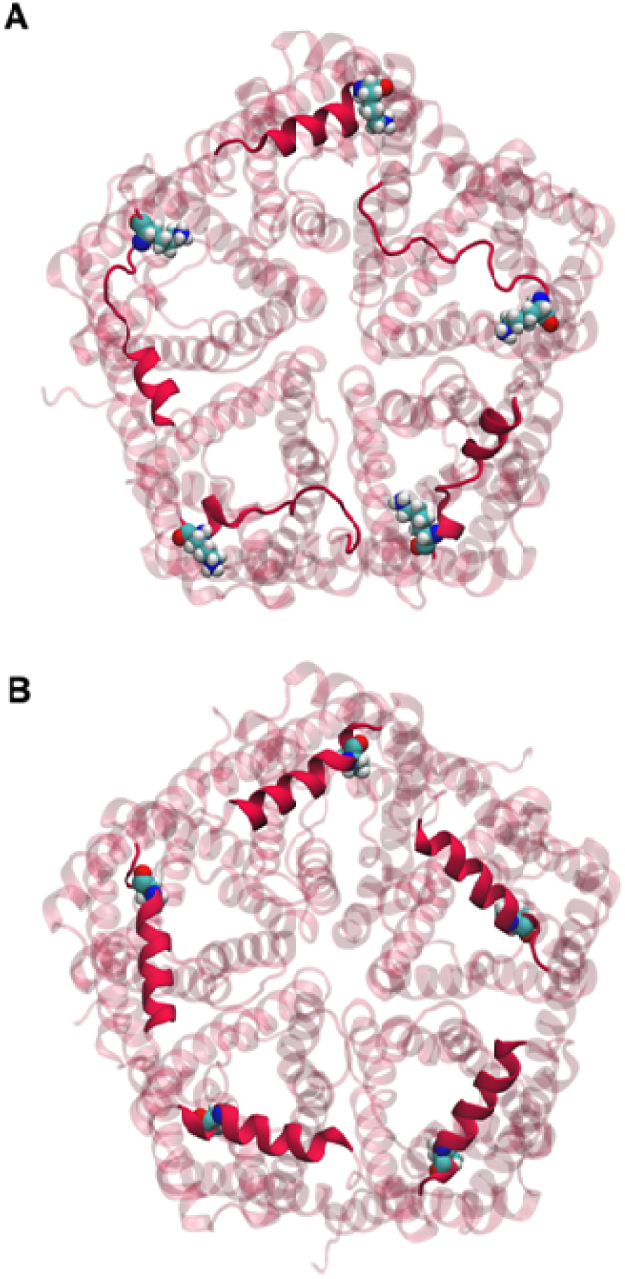
Stability of periplasmic helix (P-helix) region in EcYfdC systems. P-helix stability shown for the (A) the original homology model and (B) the revised homology model of EcYfdC channel after 500 ns production run. In the original homology model, a Lys residue faced the periplasmic entrance while in the revised model, a Val residue is at the periplasmic entrance. The P-helix is shown as opaque while the rest of the protein is shown as transparent. The position of the Lys/Val residue occupying the equivalent Lys-156 position is shown in space-filling representation.

## Potential of Mean Force profiles of formate ion and formic acid for EcYfdC channel

### Formate ion as the substrate

We used umbrella sampling technique to evaluate the energetics of substrate permeation through EcYfdC channel. PMF profiles were obtained as described in the Methods section by varying the substrates and/or the protonation state of the central His residue. Since performing the umbrella sampling on EcYfdC pentamer system is computationally expensive, we used monomers in the membrane-embedded state. A similar strategy was used in the case of VcFocA system and Lv et al. showed that VcFocA monomer in lipid bilayers remained stable [18]. They calculated the PMF profiles of formate ion and formic acid in VcFocA monomer system using adaptive biasing force calculations. We first examined the stability of EcYfdC monomer by performing 50 ns unrestrained equilibrium MD simulations. Behavior of transmembrane helical bundle, channel characteristics and the nature of two constriction sites were found be similar to that observed for the EcYfdC pentameric system (data not shown). The overall RMSD of transmembrane domain remained stable and was within 1.5 Ǻ from the starting structure. The EcYfdC monomer structure simulated at the end of this 50 ns run was used to generate starting configurations for the subsequent umbrella sampling runs.

Umbrella sampling of formate ion permeation in the EcYfdC channel was performed with the protonated central His residue. The calculated PMF profile showed several features including very high energy barriers at the periplasmic vestibule (∼35 kJ/mol) and the central constriction region (∼43 kJ/mol) (Fig 8). Inside the channel, we observed an energy minimum near the protonated central His residue (∼0.4 nm of the channel coordinate). Another energy barrier of ∼34 kJ/mol was found at the cytoplasmic slit region. PMF profiles of FocA, NirC and HSC channels have been reported in earlier biased MD studies [18,19]. The features observed in these profiles show significant differences especially at the periplasmic vestibule compared to EcYfdC channel. There is no energy barrier near the periplasmic vestibular region due to the presence of Lys residue in FocA, NirC and HSC channels. The energy minima in this region allowed the anion substrate to enter inside the channel and then the substrate moves up to the protonated central His. Due to the favorable interactions between the anionic substrate and the protonated His, the formate ion is trapped near the center of the pore. However in EcYfdC channel, due to the high energy barrier right near the periplasmic vestibule, the formate molecules do not even enter the channel. It should be noted that energy barriers exist near the constriction regions in all the channels in which the central His was considered protonated.

**Fig 8:**
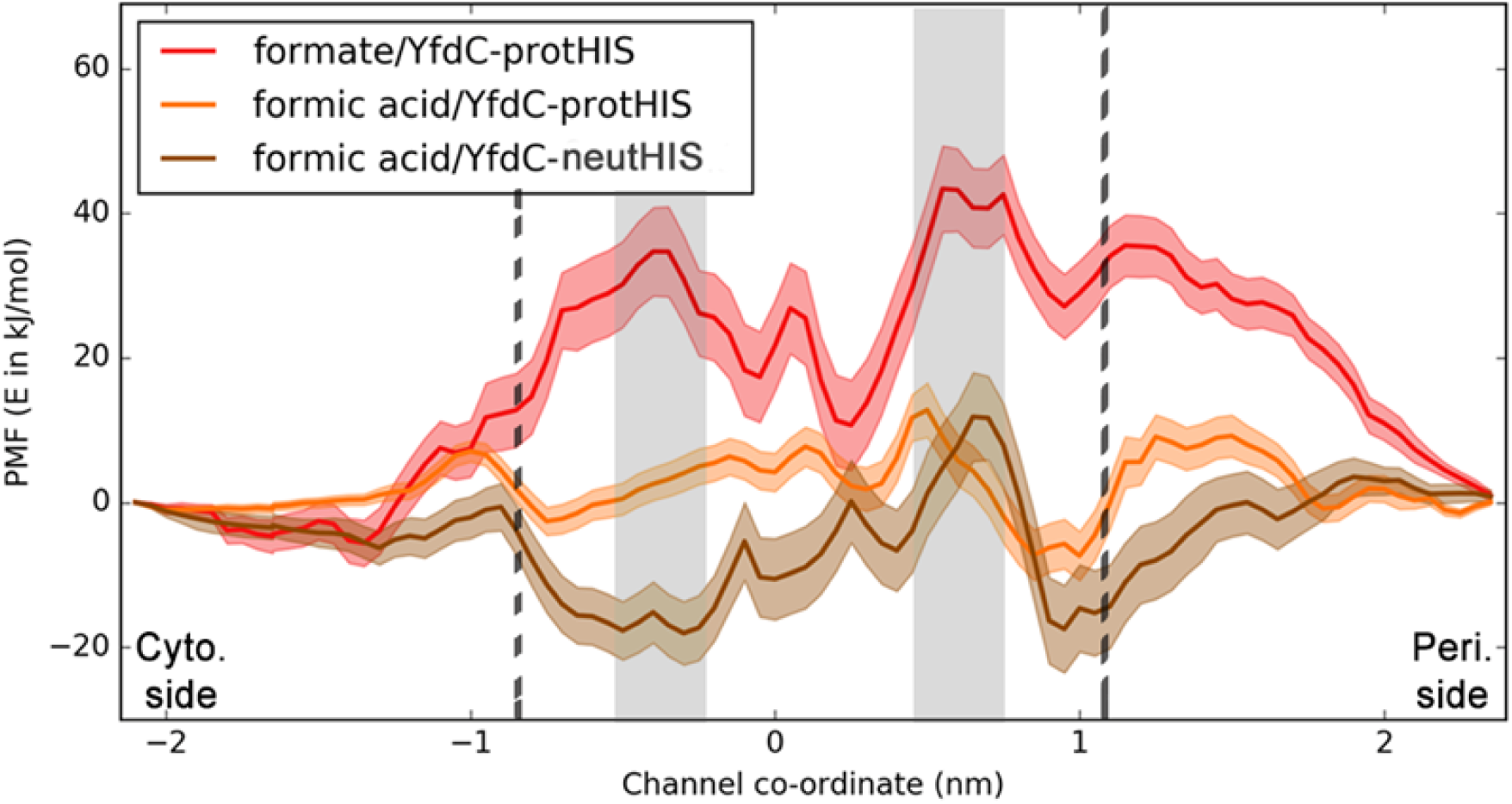
Potential of mean force (PMF) profiles of formate ion or formic acid substrates through EcYfdC channel. PMF profiles shown were calculated for substrate permeation in EcYfdC channel and the different colors correspond to different simulation conditions (Red: substrate-formate ion, central His-protonated; Orange: Substrate-formic acid, central His-protonated; Brown: substrate-formic acid, central His-neutral). Each point in the profile is shown with the standard deviation. The gray bands depict the central constriction towards the periplasmic side and the cytoplasmic slit. Locations of Glu residues at the vestibular regions are shown by dotted lines.

The energy barrier of 30 to 35 kJ/mol near the periplasmic vestibule (channel coordinate between 1.0 to 1.5 nm) is very high for the anionic substrate to enter. One reason is the absence of the highly conserved Lys residue in the periplasmic helix which is absent in YfdC subfamily. We looked at the other factors that could have contributed to this high energy barrier. We located a pore-facing Glu residue in TM5b (Glu-237 in EcYfdC) in the periplasmic vestibule (Fig 9A). The equivalent residue in other non-YfdC FNT members is Met-216 (PDB ID: 3KCU). The absence of a positively charged Lys and the presence of negatively charged Glu together are responsible for the high energy barrier for the formate anion at the periplasmic region. Similarly, another Glu residue (Glu-115 in EcYfdC; Ser-92 in VcFocA – PDB ID: 3KCU numbering) is located near the cytoplasmic side close to the cytoplasmic vestibule at ∼0.8 nm explaining high energy barrier at the opposite end (Fig 9A). Thus the two Glu residues in EcYfdC, both located in the hemi-helices outside the two constriction regions, one towards the periplasmic side and the other towards the cytoplasmic vestibule, are highly conserved (Fig 6 and Fig 9B) and seem to have a major role in the selectivity and transport mechanism in YfdC channels. To find out the significance of these two negatively charged residues and the absence of Lys at the P-helix, we performed the same calculations with formic acid, a neutral solute in EcYfdC monomer system.

**Fig 9:**
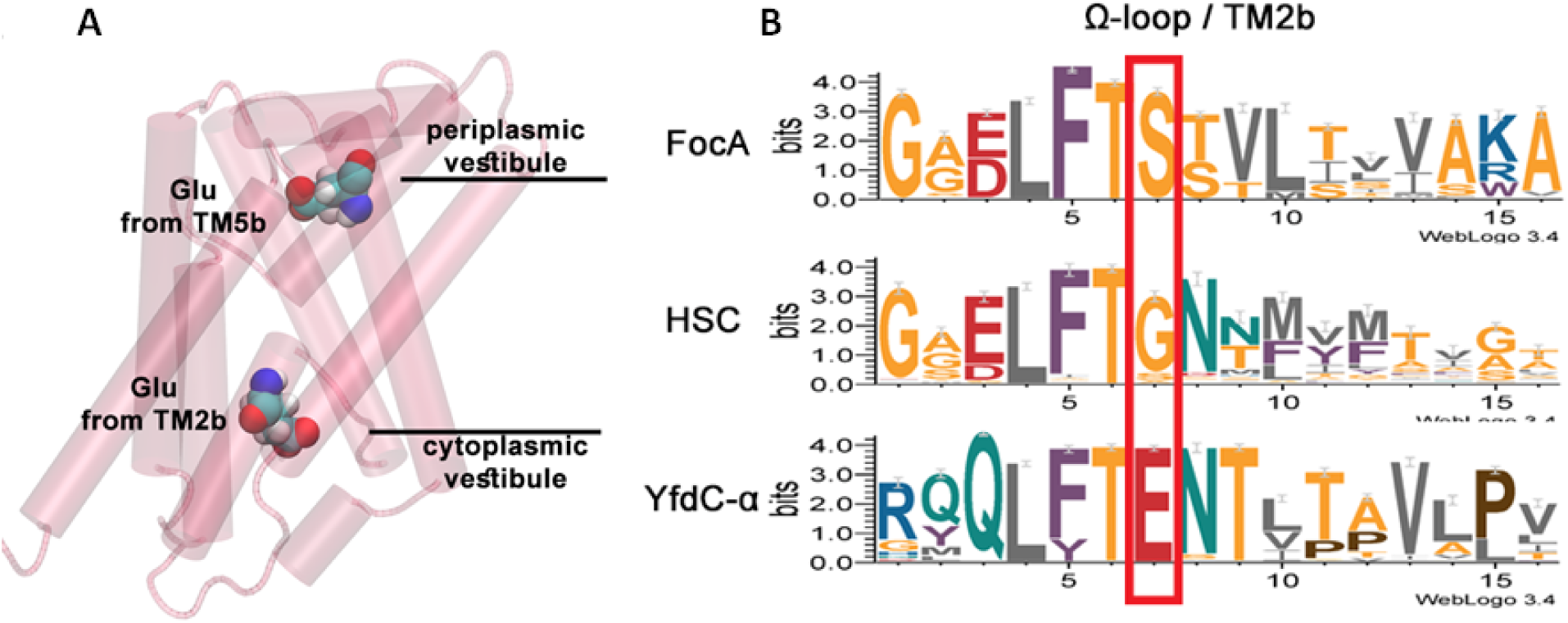
Glu residues at the channel entrance in EcYfdC. (A) Location of two Glu residues in the vestibular regions shown for the EcYfdC channel. Glu-237 and Glu-115 residues are located in TM5b and TM2b and correspond to Met-216 and Ser-92 respectively in PDB ID 3KCU numbering. (B) Sequence logo showing the conservation of Glu-115 in YfdC-α while this is substituted by small residues in FocA and HSC subfamilies.

### Formate ion versus formic acid

We calculated the energetic profile of formic acid permeation in EcYfdC channel with protonated central His. It is immediately clear that there are significant differences between the PMF profiles of formic acid and formate ions (Fig 8). The high energy barrier observed for formate ion near the periplasmic vestibule region is replaced by a smaller energy maximum for the neutral formic acid. The most notable difference is the two energy minima found near the periplasmic and cytoplasmic regions just inside the channel. This occurs at ∼ 1.0 nm distance from the center of the pore, one towards the periplasmic side and the other close to the cytoplasmic side. The energy minimum at the periplasmic side is especially pronounced and this is followed by an energy barrier near the central constriction region. As a result, a total of of nearly 20 kJ/mol energy has to be surmounted by the substrate to cross the central constriction site. This barrier is high and as a result formic acid permeation may not be feasible under physiological conditions. However, in the case of VcFocA, it has been shown that the energy barrier for formic acid is less than 8 kJ/mol when the central His is protonated [18]. This again highlights that transport properties of EcYfdC channel are different from that of typical FNT channel. The pronounced energy mimimum found at the periplasmic side could be attributed to the hydrogen bond formed between the formic acid and the side-chain carboxyl groups of Glu-237 (S2 Fig). The location of the energy minimum is roughly at the hydrogen bonding distance from the center of mass of two side-chain carboxyl oxygen atoms of Glu-237 residue.

We performed the same calculation with formic acid as the substrate and the permeation studies were performed using EcYfdC system with central His in the neutral state. The PMF profile for formic acid is very similar to that obtained for EcYfdC with protonated central His (Fig 8). However, the energy minima at both periplasmic and cytoplasmic vestibular regions are deeper indicating that the interactions between the formic acid substrate and the Glu residues at the two ends are stronger. The energy wells at both sides of the channel were located near −19.5 to −20.0 kJ/mol compared to −5.0 to −10.0 kJ/mol for EcYfdC system with protonated His. Our results indicate that EcYfdC is more amenable to permeation for neutral solutes compared to anionic substrates. Entry of formate ions followed by its protonation either at the periplasmic vestibule or near the central His region seems unlikely for EcYfdC system based on our PMF calculations and equilibraium MD studies. As a neutral solute, formic acid can easily enter the periplasmic vestibule. However, it has to overcome ∼20 kJ/mol energy barrier indicating that the rate of permeation will be low under physiological conditions.

## Glu conservation at the vestibular regions specific to YfdC-α: Clue for substrate preference?

The two Glu residues at the periplasmic and cytoplasmic vestibule regions seem to greatly influence the PMF profiles of the permeating substrates in EcYfdC channel. Glu-115 (Ser-92 in PDB ID: 3KCU numbering) is present in the TM2b half-helix close to the cytoplasmic vestibule and the second Glu residue, Glu-237 (Met-216 in PDB ID: 3KCU numbering) is found in the other half-helix TM5b near the periplasmic vestibule of the EcYfdC channel. Sequence logos of Ω-loop/TM2b and S-loop/TM5b are shown for the FocA, HSC and YfdC-α families in Fig 6 and Fig 9B respectively. Both Glu-115 and Glu-237 are more than 95% conserved in YfdC-α subfamily. It is interesting to note that Glu-115 is substituted by small residues such as Gly/Ser and Glu-237 is replaced mostly by hydrophobic residues in FocA and HSC families. The presence of negatively charged Glu residues in the vestibular regions would imply that the overall electrostatic potential in these regions also would be negative. This is evident from the calculated electrostatic potential of an EcYfdC monomer using the Adaptive Poisson-Boltzmann solver (S3 Fig). Due to the absence of the positively charged Lys residue in the periplasmic vestibule, the negative electrostatic potential has become dominant especially in the periplasmic region of EcYfdC. The same could be observed in the cytoplasmic vestibule also although to a lesser extent. It should be noted that we have not included the N-terminal region in the EcYfdC model which is likely to influence the overall electrostatic potential of cytoplasmic vestibule.

The electrostatic potential of FNT members like FocA and PfFNT (FNT homolog found in *Plasmodium falciparum*) are relatively more electropositive in the periplasmic vestibule region in contrast to that of EcYfdC channel. Weichert and Beitz have proposed that the residue Lys-156 (PDB ID: 3KCU numbering) which mainly contributes to this electropositive potential could be responsible for the protonation of the anionic substrates [20]. According to their hypothesis, the dielectric shift that causes the transfer of proton from the bulk water to the formate ion enables the substrate to permeate the channel. Recent reports also highlighted that FNT homologs in which central His is substituted by Asn or Gln also seem to exhibit the same selectivity indicating the important role of Lys-156 in protonating the substrate [29]. In the absence of Lys-156, such a possibility does not exist in EcYfdC.

## EcYfdC is not an anion channel

The above analysis indicates the reasons for a huge energy barrier for EcYfdC to transport monovalent anions. This clearly implies that unlike other FNT channel members EcYfdC is perhaps not an anion channel. Then what is its substrate selectivity? The absence of conserved Lys and the presence of conserved Glu residues at the vestibule regions indicate that it could possibly transport neutral solutes or may be cations. In this connection, two classes of membrane channels lend support to this hypothesis. The ammonium transporters AmtB transport ammonia and ammonium ion [21,22,30–32] and several aquaporins have been shown to permeate neutral solutes including ammonia [30,33,34]. Wiechert and Beitz performed experiments on *Plasmodium* FNT and the *E.coli* FocA, both are known to be selective for monovalent anions [20]. Based on their studies, they suggested that the ammonium transporters and the FNTs use similar strategy for substrate selectivity. They have used the following points to justify the claim. While the electrostatic potential of the substrate entry point in ammonium transporter is negative, FNTs exhibit a positive electrostatic field at the mouth of the channel. In both cases, the substrates have to pass through narrow hydrophobic constriction regions at the center of the channel and highly conserved histidine residue(s) present in this path have important role in the transport of solutes. Since the narrow region is highly hydrophobic, the charged species either gain or lose a proton to become neutral depending upon whether the substrate is anion or cation. By this way, they can permeate the hydrophobic region. Considering the above points, one can easily see that the characteristics of the EcYfdC channel show similarities to the ammonium transporters. Having negatively charged vestibules, a narrow hydrophobic region in the middle of the channel and a very high energy barrier at the entry point of the channel for the formate ion point to the possibility that EcYfdC, although belong to the larger FNT family, is likely to transport either neutral solutes or possibly even cationic substrates. To investigate the possibility of EcYfdC transporting neutral or cationic species, we performed additional umbrella sampling calculations using ammonia and ammonium ions as substrates for EcYfdC channel. The choice of the substrates was simply due to the fact that the ammonium transporter AmtB shares some of the characteristics described above and the structurally similar aquaporins are known to transport ammonia/ammonium ion. The substrate transported by the EcYfdC channel under physiological conditions is likely to be a different neutral or cationic species.

## PMF profiles of ammonium ion for EcYfdC channel

We performed umbrella sampling studies to calculate the PMF profiles of ammonium ion (NH_4_^+^) permeation in EcYfdC channel and followed the same protocol described earlier for determining the energetic profiles of formate ion and formic acid. We considered both protonated and neutral forms of central His and the PMF profiles calculated for both systems are shown in Fig 10A and Fig 10B. The channel coordinates ±1 nm define the vestibule regions and it is clear that the entry of ammonium ion is highly favorable in both vestibular regions irrespective of the protonation state of the central His residue. The energy well is more pronounced in EcYfdC when the central His is neutral, perhaps due to the reduced hydration within the pore (see above; Table 1). However at the center of the pore, there is a substantially high energy barrier (58 kJ/mol if the central His is protonated or 65 kJ/mol if it is neutral). This implies that the cationic ammonium ion can enter the vestibular regions but it will have difficulty crossing the central hydrophobic constriction regions due to the high energy barrier. The energy barrier could be surmounted by neutralization of the ammonium ion, as suggested for the AmtB transporters [21,31].

**Fig 10:**
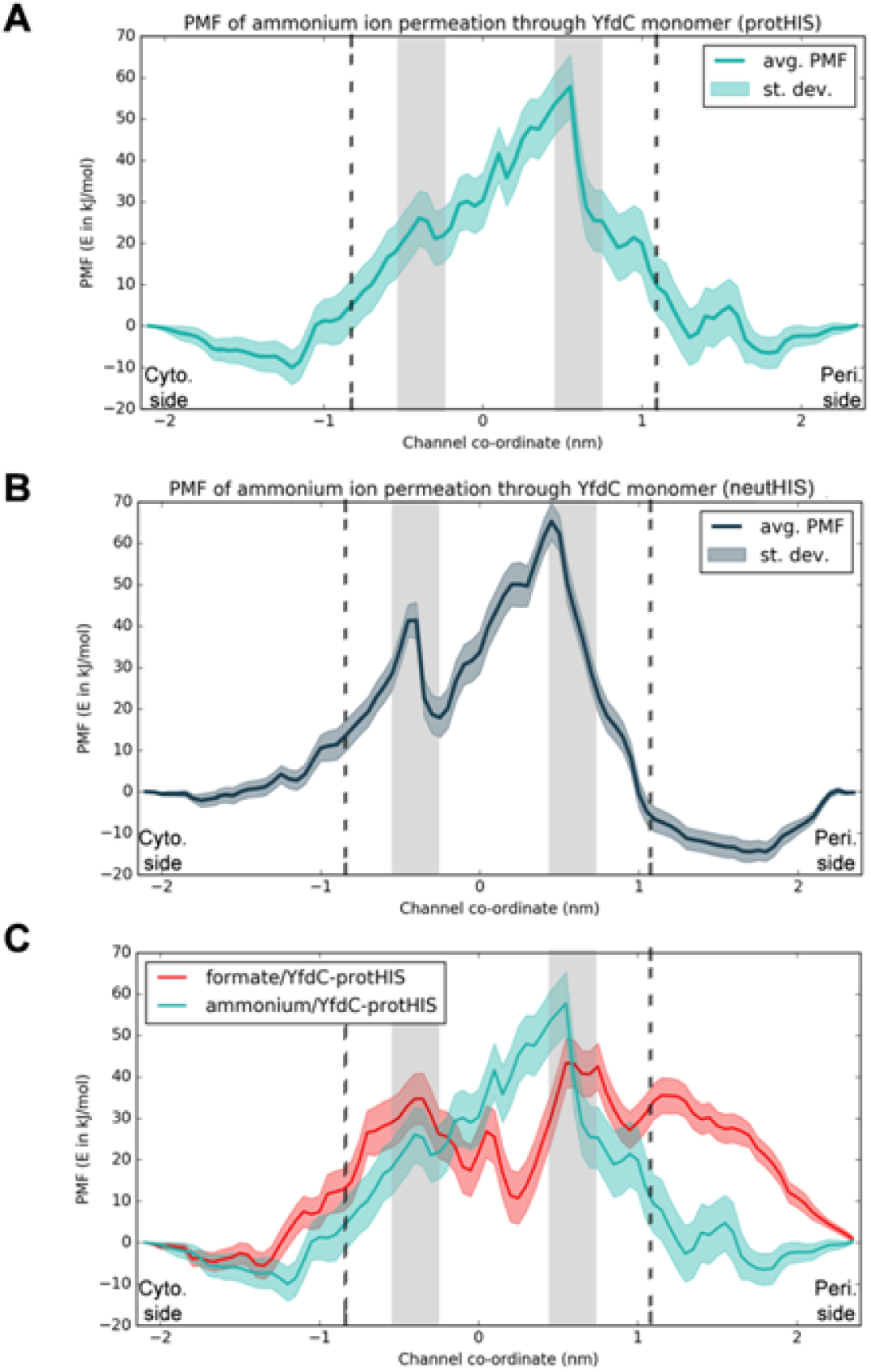
PMF profiles calculated for the permeation of ammonium ion through EcYfdC channel. PMF profiles for the permeation of ammonium ions when the central His was considered (A) in protonated and (B) in neutral form for the EcYfdC channel. (C) Comparison of PMF profiles of formate ion and ammonim ion permeation through EcYfdC channel with central His protonated. The two gray bands represent the positions of central constriction and cytoplasmic slit. Location of two conserved Glu residues at the cytoplasmic and periplasmic vestibule regions are shown in dotted lines.

It would be interesting to compare the PMF profiles of cationic ammonium ion and anionic formate ion (Fig 10C). The most striking difference is observed in the periplasmic vestibule in which formate ion has a large energy barrier even to enter inside the channel. Although the vestibule region at the cytoplasmic side shows difference, it is subtle and it could be due to the fact that we did not include the N-terminal helical region in our calculations (see above).

## PMF profiles of ammonia for the EcYfdC channel

We repeated the above calculations to evaluate the PMF profiles of ammonia as substrate for the EcYfdC channel by considering the central His in both protonated and neutral form. PMF profiles of neutral ammonia revealed that there is a small energy barrier of 5 to 10 kJ/mol at the vestibular regions (Fig 11A and Fig 11B). This can be easily understood as the Glu residue in the vestibule cannot form strong interactions with the passing solute indicating that the substrate will not be trapped in an energy well. This is in contrast to what was found for formic acid whose energy profile showed a deep energy well due to the hydrogen bond formation between the formic acid and the Glu residue at the periplasmic vestibule creating an energy barrier of close to 35 kJ/mol (Fig 8).

**Fig 11:**
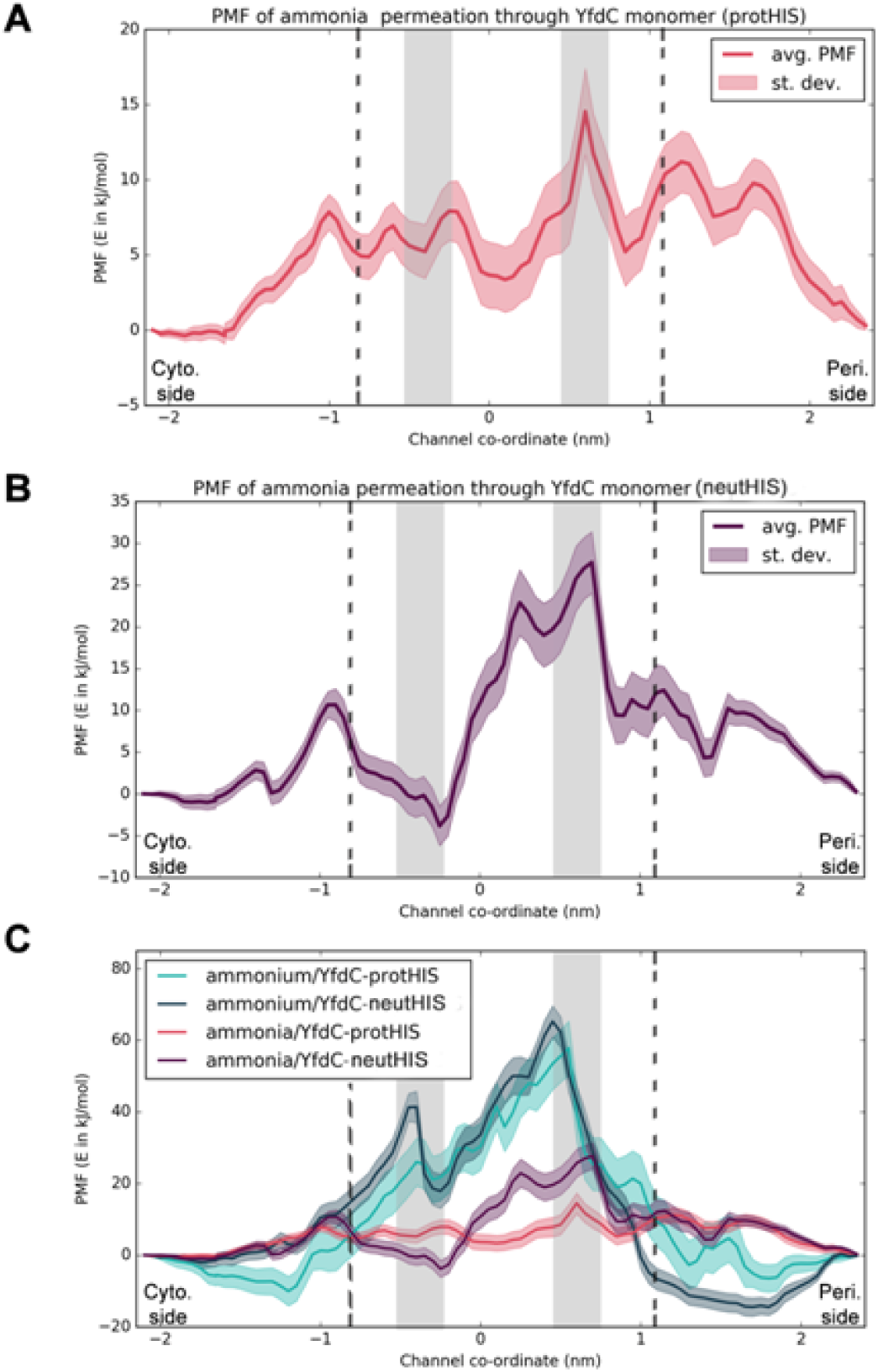
PMF profiles of ammonia permeation through EcYfdC channel. Ammonia PMF profiles calculated with the central His (A) in protonated and (B) in neutral forms. (C) Comparison of PMF profiles of ammonium ion and ammonia permeation through EcYfdC channel with central His considered in both protonated and neutral forms. The two gray bands depict the location of cytoplasmic slit and the central constriction region. The dotted lines represent the positions of the two conserved Glu resides towards the periplasmic and cytoplasmic vestibule regions

Comparing the energy profiles of ammonium ion and ammonia for both protonated and neutral His residue, we could make the following observations (Fig 11C). The PMF profiles obtained for the ammonium ion look similar for both the simulations in which the central His was considered protonated or neutral. However for the neutral ammonia, there are some notable differences. The central constriction exhibits an energy barrier of ∼14.5 kJ/mol when the His is protonated. For the system with neutral His, the energy profile in the same region shows a higher barrier of close to 28 kJ/mol implying that ammonia will have lower probability of permeating EcYfdC if the central His is neutral. Neutral His in the central constriction makes this region more hydrophobic resulting in increased height of the energy barrier. Hence, complete permeation of ammonia is less probable in EcYfdC with the neutral His as found for the formic acid substrate (Fig 8). However the energy barrier of 14 kJ/mol for the EcYfdC system with proptonated His is very similar to that observed for bacterial GlpF channel and human AQP4 [33,35], both are members of MIP superfamily. A slightly higher energy barrier is observed for human AQP1 and also for the ammonia gas permeating through POPE bilayer [35]. As MIP members with similar fold have been shown to permeate ammonia in oocyte membranes, it is tempting to speculate that EcYfdC channel, when the central His is protonated, is likely to transport neutral solutes similar to ammonia.

## Conclusion

In a recent bioinformatics analysis of FNT family members, 27% of the identified FNTs were classified as members belonging to YfdC subfamily and they were further subdivided into two groups, YfdC-α and YfdC-β. YfdC subfamily members are functionally uncharacterized and this amounts to nearly one third of all FNT members. In this study, we have investigated the possible function of *E. coli* YfdC channel using computational techniques. We first performed equilibrium MD simulation of EcYfdC in presence of formate ion and formic acid substrates and compared the simulation results with that of VcFocA and CdHSC channels. Our results clearly demonstrated that the anionic substrates do not enter the vestibule regions of EcYfdC while they crossed the vestibule regions and interacted strongly with the central His residue in VcFocA and CdHSC channels. While a Lys residue in the periplasmic helix and its interacting partners are highly conserved in FocA, HSC and NirC subfamilies, they are conspicuously absent in YfdC subfamily members. Similarly while the YfdC-α members have nearly invariant Glu residues at the vestibule regions, they are noticeably absent in non-YfdC members. This contrasting conservation pattern in the vestibule regions results in electrostatic potential that is predominantly negative in YfdC-α members and it is quite opposite in non-YfdC members of FNT family which have electropositive potential in the periplasmic vestibule. This explains why anionic substrates cannot enter the vestibule regions of EcYfdC raising a doubt whether EcYfdC is an anion-specific channel. It is quite possible that EcYfdC transports neutral molecules, or perhaps even cationic species.

To identify the substrate preference of EcYfdC channel, we performed umbrella sampling and evaluated the PMF profiles of formate anion, neutral formic acid and ammonia and cationic ammonium ion. Comparison of PMF profiles of formate anion and the neutral formic acid immediately revealed the absence of the huge energy barrier for the formic acid substrate at the periplasmic vestibule region. While formate anion entry inside EcYfdC channel is ruled out, our results indicate that formic acid is likely to encounter a 20 kJ/mol energy barrier. Hence the permeation rate of formic acid is likely to be low. Although the overall fold is completely different, ammonium transporters and YfdC subfamily members share certain features. This includes a negative electrostatic potential at the vestibule regions, a narrow hydrophobic constriction in the middle of the channel and the presence of one or more highly conserved His residues near the hydrophobic constriction. Although there is no experimental evidence that ammonia/ammonium ion is likely to be the endogenous substrate for EcYfdC channel, we evaluated the PMF profiles of ammonia and ammonium ion for the EcYfdC channel. Our results clearly demonstrated that the EcYfdC can transport neutral solutes like ammonia when the central His is protonated. The energy barrier of 14 kJ/mol is relatively small and comparable to the barriers observed for solutes that are transported with high efficiency by other channels. Hence, we conclude that EcYfdC channel is not an anion-transporting channel although it belongs to the anion-transporting FNT family. The pattern of YfdC-specific sequence conservation and our calculations clearly illustrate that EcYdC channel can transport neutral solutes or even cations. This is most likely applicable to other members of YfdC subgroup. This is one more example of proteins adopting the same fold to perform different functions. Experimental studies can be carried out to find out whether EcYfdC can transport ammonia and the results of such studies will strengthen the conclusions arrived at this study.

## Materials and Methods

Three representative members of the FNT family, formate channel from *V. cholera* (VcFocA), hydrosulphide channel from *Clostridium difficile* (CdHSC) and the uncharacterized YfdC channel from *E. coli* (EcYfdC) were selected for all-atom molecular dynamics simulations. Their UniProt [36] IDs are Q9KRE7, Q186B7 and P37327 respectively. Three-dimensional structures of VcFocA and CdHSC have been determined experimentally and were downloaded from the Protein Data Bank (PDB) [37]. The structures corresponding to the PDB IDs 3KLY (resolution: 2.1 Ǻ) [12] and 3TE0 (resolution: 2.09 Ǻ) [7] served as starting structures for VcFocA and CdHSC channels respectively. All heteroatoms were removed from the PDB structures and the missing residues were built using UCSF Chimera suite [38]. In the case of CdHSC structure, the K148E mutation was reversed *in silico* to the wild-type residue. Since the region corresponding to N-terminal helix is disordered in VcFocA structure, this N-terminal helical region was removed in CdHSC structure. The role of N-terminal helix in the substrate permeation has not been clearly established in previous studies [18].

Members of YfdC family are not structurally and functionally characterized. With no structural information available, we constructed a three-dimensional model for the EcYfdC using a protocol similar to that we used previously to model FNTs, MIPs and other membrane proteins [9,13,39,40]. Homology modeling using multi-template approach as implemented in MODELLER v9.13 [41] was used to build the EcYfdC model. Structures of EcFocA (PDB ID: 3KCU [11]; Chain B), VcFocA (PDB ID: 3KLY [12]; Chain E), CdHSC (PDB ID: 3TE0 [7]; Chain E) and StNirC (PDB ID: 4FC4 [4]; Chain I) were chosen as templates to represent the three FNT subfamilies. The Ω-loop and TM2b regions were modeled as helical based on CdHSC and StNirC structures. We did not include N-terminal region for modeling. Among the 100 models generated, we selected the best model based on the optimum score derived from MODELLER’s objective function. Side-chains were refined using the graph theory-based SCRWL4 method [42]. The model thus generated was energy minimized using GROMACS ver 5.0.6 [43] with CHARMM36 force field [44]. This minimized model was used to generate the EcYfdC pentamer by superimposition on the CdHSC structure (PDB ID: 3TE0) and the resultant pentamer was further energy minimized.

### MD Simulations of FNT members

We used CHARMM-GUI Membrane Builder suite [45] to generate the lipid-embedded FNT pentamer system. Each pentamer representing three FNT subfamilies were inserted in a pre-equilibrated POPE (1-palmitoyl-2-oleoyl-sn-glycero-3-phosphoethanolamine) lipid bilayer patch containing ∼280 lipids. Both sides of the bilayers were solvated and adequate counter ions were included to neutralize the system. The central pore formed by the five monomers was plugged using two POPE lipids from the periplasmic side. All three proteins (VcFocA, CdHSC and EcYfdC) were simulated with saturating concentration of 120 mM solute. The solute was either formate or formic acid and the central conserved His was considered as protonated or neutral. With this combination, we have simulated nine systems using CHARMM36 force-filed [44] and all simulated systems are summarized in Table 1. We have not considered the formate substrate in combination with neutral His as such a system is highly unlikely to occur due to energetic constraints.

All simulations were performed using GROMACS v5.0.6 [43]. Each system was energy minimized followed by a seven-step NPT equilibration procedure as suggested by the CHARMM-GUI Membrane Builder Suite [45]. Positional restraints on protein heavy atoms and lipid headgroups and dihedral restraints on protein side-chains were applied and were gradually released in six steps over a period of 375 ps. Berendesen’s temperature and pressure coupling methods [46] were applied during this process. A final 20 ns equilibration was performed free of all the restraints to make sure that the systems are relaxed completely. The reference pressure was 1 bar and was maintained using a semi-isotropic pressure coupling to a Parrinello-Rahman pressure coupling barostat [47]. The reference temperature of 310 K was maintained using Nose-Hoover coupling algorithm [48]. Long-range interactions were evaluated using Particle-Mesh Ewald (PME) method with Verlet cut-off scheme. A cut-off of 1.2 nm was used to calculate the Coulombic and VDW interactions in real space. Periodic boundary conditions were applied in all directions. After equilibration, a production run of 500 ns was performed on each of the nine systems considered in this study. MD simulated structures were saved at 10 ps interval for analysis.

### Potential of Mean Force (PMF) profiles for substrate permeation

Umbrella sampling technique [49] was used on FNT channel members to obtain the potential of mean force (PMF) profiles of permeation of various substrates. In previous studies, adaptive biasing force calculations were performed on VcFocA monomer rather than a pentamer to minimize the computational cost [18]. In the present study, we have adopted a similar approach to obtain PMF profiles and the studies were carried out on VcFocA and EcYfdC monomeric systems. The preparation of the systems for simulations was similar to that followed for the pentameric systems (see above). The monomer (EcYfdC or VcFocA) was first embedded in a POPE bilayer and solvated using TIP3P waters [50] with added counter-ions. CHARMM-GUI Membrane Builder [45] was used for this purpose. Equilibration of the monomer system was carried out in seven stages very similar to that adopted for the pentameric system. At the last stage, we have performed a longer equilibration of 15 ns without any restraints and the structure generated from this procedure served as the starting configurations for the umbrella sampling studies. All simulation parameters were same as that used in the equilibrium simulation studies of pentameric systems.

We used Steered Molecular Dynamics (SMD) as implemented in GROMACS v5.0.6 to generate the starting configurations for umbrella sampling. The reaction coordinates were defined as the distance along the Z-axis, perpendicular to the membrane plane. Distance is calculated between the center of mass of the substrate and the center of mass of the Cα atoms of the transmembrane region. Substrate was initially placed at the cytoplasmic vestibule and steered gradually along the Z-axis towards the extracellular side. The pulling was performed 0.5 nm/ns with a force constant of 5000 kJ/mol/nm. In this way, 90 starting points were selected along the reaction pathway and the separation between the umbrellas was 0.05 nm. At each umbrella window, 1 ns equilibration was performed with the positions of the substrate atoms restrained with a force constant of 10,000 kJ/mol/nm. This was followed by a further 1.5 ns simulation for each umbrella window. Harmonic restraints were applied to hold the substrate at the center of the umbrella with force constant up to 2500 kJ/mol/nm. Flat-bottomed potential was employed on the substrate molecule to restrict it within a cylinder of radius 0.5 nm. We also restrained counter-ions in the bulk-water slabs using flat-bottomed potentials. Phosphorous atoms of lipid head-groups which were farther than 0.5 nm were also held along the Z-axis to prevent the protein from moving along the axis normal to the membrane plane during the umbrella samplings. To enhance the confidence of the PMF calculations, we repeated the umbrella simulations five times using different initial velocities for each umbrella run. Thus each PMF calculation is the result of over one microsecond simulation with the PMF profiles being calculated from 700 ns of production run.

Average PMF profiles were calculated using the periodic weighted histogram analysis (WHAM) method [51] with g_wham application [52] as implemented in GROMACS v5.0.6. Statistical errors were estimated using 20 Bayesian bootstraps. Trapezoidal correction was applied to the vestibular ends of all the calculated PMFs using the theory as described in previous studies [19,35]. In addition to formate and formic ion, ammonia and ammonium ions were also used as substrates for the umbrella sampling studies performed on EcYfdC. CHARMM36 Force-filed parameters are available for all the substrates except ammonium ion. Parameters for ammonium ion were derived using the method described by Nygaard et al. [53].

### Temporal channel radius profiles

Channel radius profiles of all monomers were evaluated as a function of time using the program HOLE [24]. We studied the evolution of average channel radii at each point along the channel axis over the course of 500 ns simulation time and plotted a two-dimensional profile for each simulated system. We have previously used temporal channel radius profiles in understanding the cooperativity between monomers of plant aquaporin tetramers [25]. In the present study, temporal radius profiles of FNT members were used to monitor the behavior of two constriction regions and compared them in the presence of different substrates with central His protonated or neutral. In a pentameric system, channel radius profile was calculated for every monomer at 10 ps intervals. Data from all five monomers for every 2 ns simulation window was averaged and the average channel radius profiles for 250 simulation windows were used to plot the two-dimensional temporal channel radius profiles for each simulated FNT system.

### Substrate and solvent permeation studies

We monitored all the permeation events of the substrate moieties or solvent molecules. A permeation event is said to happen when there is a complete passage of the substrate/solvent through a cylinder of 0.5 nm radius passing through the center of each monomer along the Z-axis. Sporadic or partial permeation events along the central pore or the lipid bilayer as seen in the case of neutral substrates were ignored. Permeation events were monitored using the in-house python scripts. The dipole orientation of permeating solvent molecules was determined using the gmx_h2order module as available in GRAMACS package.

## Acknowledgements

High Performance Computing facility of IIT-Kanpur is gratefully acknowledged. RS is Pradeep Sindhu Chair Professor at IIT-Kanpur. MM thanks IIT-Kanpur for a fellowship. We thank all our lab members for useful discussions.

## Supporting Information

**S1 Fig: Temporal channel radius profile of the revised EcYfdC model and the MD trajectories of Center of mass Z-coordinates of the formate ions along the channel coordinate**

(A) Temporal channel radius profile for the revised EcYfdC model plotted for the entire 500 ns production run. (B) Z-coordinates of the center-of-mass of individual formate ions are plotted along the channel coordinate for the 500 ns production run. Each color depicts one unique formate ion substrate. Formate ions did not enter the vestibule regions of EcYfdC. Position ‘0’ in Y-axis corresponds to the central His residue. The positive and negative Y-axis corresponds to the periplasmic and cytoplasmic sides respectively.

**S2 Fig: Favorable interactions of formic acid substrate in the periplasmic vestibule**

The substrate formic acid shown in space-filling representation participates in favorable hydrogen bond interactions with the conserved periplasmic Glu residue in TM5b segment (Glu-237 in PDB ID: 3KCU numbering). The central His residue and the conserved Glu are shown in stick representation.

**S3 Fig: Electrostatic potential map of EcYfdC protein**

Electrostatic potential surface map calculated using Adaptive Poisson-Boltzmann Solver for the EcYfdC protein. The electronegative potential at the periplasmic vestibule is prominent. Cytoplasmic vestibule also shows electronegative surface due to another conserved Glu residue. It should be noted that the N-terminal segment of EcYfdC which is present in the cytoplasmic side was not included in the simulations due to differences in the structures of different FNT homologs.

## Data Availability

All the relevant data are provided within the manuscript and its Supporting Information files.

## References

1. Marchetti RV, Lehane AM, Shafik SH, Winterberg M, Martin RE, et al. (2015) A lactate and formate transporter in the intraerythrocytic malaria parasite, *Plasmodium falciparum*. Nature Commn 6: Art. No. 6721.

2. Wu B, Rambow J, Bock S, Holm-Bertelsen J, Wiechert M, et al. (2015) Identity of a *Plasmodium* lactate/H+ symporter structurally unrelated to human transporters. Nature Commn 6.

3. Lu W, Schwarzer NJ, Wacker T, Andrade SLA, Einsle O (2013) The formate/nitrite transporter family of anion channels. Biol Chem 394: 715–727.

4. Lu W, Du J, Schwarzer NJ, Gerbig-Smentek E, Einsle O, et al. (2012) The formate channel FocA exports the products of mixed-acid fermentation. Proc Natl Acad Sci USA 109: 13254–13259.

5. Waight AB, Czyzewski BK, Wang D-N (2013) Ion selectivity and gating mechanisms of FNT channels. Curr Opin Struct Biol 23: 499–506.

6. Clegg S, Yu F, Griffiths L, Cole JA (2002) The roles of the polytotic membrane proteins Nark, Naru and NirC in Escherichia coli K-12: two nitrate and three nitrite transporters. Mol Microbiol 44: 143–155.

7. Czyzewski BK, Wang D-N (2012) Identification and characerization of a bacterial hydrosulphide channel. Nature 483: 494–497.

8. Wiechert M, Erler H, Golldack A, Beitz E (2017) A widened substrate selectivity filter of eukaryotic formate-nitrite transporters enables high level lactate conductance. FEBS J 284: 2663–2673.

9. Mukherjee M, Vajpai M, Sankararamakrishnan R (2017) Anion-selective formate/nitrite transporters: taxonomic distribution, phylogenetic analysis and subfamily-specific conservation pattern in prokaryotes. BMC Genomics 18: Art. No. 560.

10. Suppmann B, Sawers RG (1994) Isolation and characterization of hypophosphite-resistant mutants of *Escherichia coli*: identification of FocA protein, encoded by the *Pfl* operon, as a putative formate transporter. Mol Microbiol 11: 965–982.

11. Wang Y, Huang Y, Wang J, Cheng C, Huang W, et al. (2009) Structure of the formate transporter FocA reveals a pentameric aquaporin-like channel. Nature 462: 467–472.

12. Waight AB, Love J, Wang D-N (2010) Structure and mechanism of pentameric formate channel. Nature Struct Mol Biol 17: 31–37.

13. Verma RK, Gupta AB, Sankararamakrishnan R (2015) Major intrinsic protein superfamily: Channels with unique structural features and diverse selectivity filters. Methods Enzymol 557: 485–520.

14. Hunger D, Doberenz C, Sawers RG (2014) Identification of key residues in the formate channel FocA that control import and export of format. Biol Chem 395: 813–825.

15. Hunger D, Rocker M, Falke D, Lilie H, Sawers RG (2017) The C-terminal six amino acids of the FNT channel FocA are required for formate translocation but not homopentamer integrity. Front Microbiol 8: Art. No. 1616.

16. Feng Z, Hou T, Li Y (2012) Concerted movement in pH-dependent gating of FocA from molecular dynamics simulations. J Chem Inf Model 52: 2119–2131.

17. Lu W, Du J, Wacker T, Gerbig-Smentek E, Andrade SLA, et al. (2011) pH-dependent gating in a FocA formate channel. Science 332: 352–354.

18. Lv X, Liu H, Ke M, Gong H (2013) Exploring the pH-dependent substrate transport mechanism of FocA using molecular dynamics simulation. Biophys J 105: 2714–2723.

19. Atkovska K, Hub JS (2017) Energetics and mechanism of anion permeation across formate-nitrite transporters. Sci Rep 7: Art. No. 12027.

20. Wiechert M, Beitz E (2017) Mechanism of formate-nitrite transporters by dielectric shift of substrate acidity. EMBO J 36: 949–958.

21. Khademi S, O’Connell III J, Remis J, Robles-Colmenares Y, Miercke LJ, et al. (2004) Mechanism of ammonia transport by Amt/MEP.Rh: Structure of AmtB at 1.35 A. Science 305: 1587–1594.

22. Zheng L, Kostrewa D, Berneche S, Winkler FK, Li X-D (2004) The mechanism of ammonia transport based on the crystal structure of AmtB of *Escherichia coli*. Proc Natl Acad Sci USA 101: 17090–17095.

23. Doberenz C, Zorn M, Falke D, Nannemann D, Hunger D, et al. (2014) Pyruvate formate-lyase interacts directly with the formate channel FocA to regulate formate translocation. J Mol Biol 426: 2827–2839.

24. Smart OS, Neduvelil JG, Wang X, Wallace BA, Sansom MSP (1996) HOLE: A program for the analysis of the pore dimensions of ion channel structure models. J Mol Graph 14: 354–360.

25. Vajpai M, Mukherjee M, Sankararamakrishnan R (2018) Cooperativity in plant plasma membrane intrinsic proteins (PIPs): Mechanism of increased water transport in maize PIP1 channels in hetero-tetramers. Sci Rep 8: Art. No. 12055.

26. de Groot BL, Grubmuller H (2001) Water permeation across biological membranes: Mechanism and dynamics of aquaporin-1 and GlpF. Science 294: 2353–2357.

27. Tajkhorshid E, Nollert P, Jensen MO, Miercke LJ, O’Connell JO, et al. (2002) Control of the selectivity of the aquaporin water channel family by global orientational tuning. Science 296: 525–530.

28. de Groot BL, Grubmuller H (2005) The dynamics and energetics of water permeation and proton exclusion in aquaporins. Curr Opin Struct Biol 15: 176–183.

29. Helmstetter F, Arnold P, Hoger B, Petersen LM, Beitz E (2019) Formate-nitrite transporters carrying nonprotonatable amide amino acids instead of a central histidine maintain pH-dependent transport. J Biol Chem 294: 623–631.

30. Musa-Aziz R, Chen L-M, Pelletier MF, Boron WF (2009) Relative CO2/NH3 selectivities of AQP1, AQP4, AQP5, AmtB, and RhAG. Proc Natl Acad Sci USA 106: 5406–5411.

31. Javelle A, Lupo D, Ripoche P, Fulford T, Merrick M, et al. (2008) Substrate binding, deprotonation, and selectivity at the periplasmic entrance of *Escherichia coli* ammonia channel AmtB. Proc Natl Acad Sci USA 105: 5040–5045.

32. Javelle A, Lupo D, Zheng L, Li X-D, Winkler FK, et al. (2006) An unusual twin-His arrangement in the pore of ammonia channels is essential for substrate conductance. J Biol Chem 281: 39492–39498.

33. Assentoft M, Kaptan S, Schneider H-P, Deitmer JW, de Groot BL, et al. (2016) Aquaporin 4 as a NH_3_ chennel. J Biol Chem 291: 19184–19195.

34. Kirscht A, Kaptan SS, Bienert GP, Chaumont F, Nissen P, et al. (2016) Crystal structure of ammonia-permeable aquaporin. PLoS Biology 14: Art. No. e1002411.

35. Hub JS, de Groot BL (2008) Mechanism of selectivity in aquaporins and aquaglyceroporins. Proc Natl Acad Sci USA 105: 1198–1203.

36. Consortium TU (2017) UniProt: the universal protein knowledgebase. Nucleic Acids Res 45: D158–D169.

37. Berman HM, Westbrook J, Feng Z, Gilliiland G, Bhat TN, et al. (2000) The Protein Data Bank. Nucleic Acids Res 28: 235–242.

38. Pettersen EF, Goddard TD, Huang CC, Couch GS, Greenblatt DM, et al. (2004) UCSF Chimera - a visualization system for exploratory research and analysis. J Comput Chem 25: 1605–1612.

39. Gupta AB, Sankararamakrishnan R (2009) Genome-wide analysis of major intrinsic proteins in the tree plant Populus trichocarpa: Characterization of XIP subfamily of aquaporins from evolutionary perspective. BMC Plant Biol 9: Art. no. 134.

40. Gupta A, Sankararamakrishnan R (2018) dbSWEET: An integrated resource for SWEET superfamily to understand, analyze and predict the function of sugar transporters in prokaryotes and eukaryotes. J Mol Biol: doi: 10.1016/j.jmb.2018.1004.1013.

41. Webb B, Sali A (2014) Comparative protein structure modeling using MODELLER. Curr Protocols Bioinfo 47: 5.6.1–5.6.32.

42. Krivov GG, Shapovalov MV, Dunbrack J, R. L. (2009) Improved prediction of protein side-chain conformations with SCWRL4. Proteins: Struct Func Bioinf 77: 778–795.

43. Pronk S, Pall S, Schulz R, Larsson P, Bjelkmar P, et al. (2013) GROMACS 4.5: a high-throughput and highly parallel open source molecular simulation toolkit. Bioinformatics 29: 845–854.

44. Huang J, Mackerell AD Jr. (2013) CHARMM36 all-atom additive protein force field: Validation based on comparison to NMR data. J Comput Chem 34: 2135–2145.

45. Jo S, Lim JB, Klauda JB, Im W (2009) CHARMM-GUI membrane builder for mixed bilayers and its application to yeast membranes. Biophys J 97: 50–58.

46. Berendsen HJC, Postma JPM, van Gunsteren WF, DiNola A, Haak JR (1984) Molecular dynamics with coupling to an external bath. J Chem Phys 81: 3684–3690.

47. Parrinello M, Rahman A (1981) Polymorphic transitions in single crystals: A new molecular dynamics method. J Appl Phys 52: 7182–7190.

48. Evans DJ, Holian BL (1985) The Nose-Hoover thermostat. J Chem Phys 83: 4069–4074.

49. Kastner J (2011) Umbrella sampling. WIREs Comput Mol Sci 1: 932–942.

50. Jorgensen WL, Chandrasekhar J, Madura JD, Impey RW, Klein ML (1983) Comparison of simple potentil functions for simulating liquid water. J Chem Phys 79: 926–935.

51. Kumar S, Rosenberg JM, Bouzida D, Swendsen RH, Kollman PA (1992) The weighted histogram analysis method for free-energy calculations on biomolecules. I. The method. J Comput Chem 13: 1011–1021.

52. Hub JS, de Groot BL, van der Spoel D (2010) g_wham - A free weighted histogram analysis implementation including robust error and autocorrelation estimates. J Chem Theory Comput 6: 3713–3720.

53. Nygaard TP, Rovira C, Peters GH, Jensen MO (2006) Ammonium recruitment and ammonia transport by *E.coli* ammonia channel AmtB. Biophys J 91: 4401–4412.

